# Communication determines population-level fitness under cation stress by modulating the ratio of motile to sessile *B. subtilis* cells

**DOI:** 10.1101/2021.11.30.470380

**Authors:** Benedikt K. Steinfeld, Qinna Cui, Tamara Schmidt, Ilka B. Bischofs

## Abstract

Bacterial populations frequently encounter potentially lethal environmental stress factors. Growing *Bacillus subtilis* populations are comprised of a mixture of “motile” and “sessile” cells but how this affects population-level fitness under stress is poorly understood. Here, we show that, unlike sessile cells, motile cells are readily killed by monovalent cations under conditions of nutrient deprivation – owing to elevated expression of the *lytABC* operon, which codes for a cell-wall lytic complex. Forced induction of the operon in sessile cells also causes lysis. We demonstrate that population composition is regulated by the quorum sensing regulator ComA, which can favor either the motile or the sessile state. Specifically social interactions by ComX-pheromone signaling enhance population-level fitness under stress. Our study highlights the importance of characterizing population composition and cellular properties for studies of bacterial physiology and functional genomics. Our findings open new perspectives for understanding the functions of autolysins and collective behaviors that are coordinated by chemical and electrical signals, with implications for multicellular development and biotechnology.

## Introduction

Many bacterial populations frequently encounter potentially lethal environmental stress factors, and have evolved strategies that aid their survival. The Gram-positive *Bacillus subtilis* is an environmentally ubiquitous bacterium with a complex life style, and exhibits many beneficial traits that are exploited in diverse biotechnological applications. Studies of *B. subtilis* survival traits have a long history, and have contributed to a better understanding of how the species is so successful. In addition, this line of inquiry can help to identify how populations can be controlled and engineered to better serve mankind, specifically when naturally acquired resistance traits interfere with desired applications.

Vegetative *B. subtilis* can exist in two phenotypically distinct states: as a flagellated, motile cell that can explore its environment; and as a sessile cell that produces an extracellular matrix and can exploit locally available resources^1,2^. Motile cells have a short-rod like morphology while sessile cells tend to grow in chains of connected cells and both cell-types are capable of growing and dividing^3^ The transitions between the motile and a sessile state are controlled by a complex regulatory network and can be modulated by environmental and cellular signals as well as noise^4,5^. Circumstantial evidence has also linked cell-cell communication via chemical signals to coordinate the physiologies of the different cell types during biofilm formation, via the lipopeptide surfactin^6^, although this has been recently questioned^7^.

It is tempting to equate a motile cell with a “planktonic cell” and a sessile cell with a “biofilm cell”. However, there exists no planktonic or biofilm cell *per se*, since both biofilms and planktonic cultures are comprised of a mixture of cell types^4,5,8,9^. Thus, although biofilms are generally considered to be more stress resistant than planktonic cultures, it is unclear whether individual sessile cells are actually more robust than motile cells. In addition to that, an important implication of phenotypic diversification within isogenic populations is that population composition becomes a crucial determinant of population-level fitness under stress. Nevertheless, this fact has been – and often still is – neglected^10–19^.

Here we show that individual sessile and motile cells exhibit different levels of stress tolerance, and that social interactions by chemical communication shift population composition which enhances population-level fitness under stress. Notably, this study was not originally designed to address these issues. Its roots lay in the serendipitous discovery that a mutant with impaired chemical communication lysed quickly in standard phosphate-buffered saline (PBS). We then employed a combination of bulk and single-cell studies to elucidate the basis for this phenomenon, which revealed unexpected links between chemical signaling and stress tolerance, and indications of a role for electrical signaling via monovalent cations. We specifically found that the quorum sensing master regulator ComA regulates population composition and favors the motile or sessile state depending on its phosphorylation status (and independent of surfactin). Motile cells are specifically sensitized to killing by monovalent cations by elevated expression the LytABC autolysin complex. This finding raises the possibility that motile subpopulations could be specifically targeted by electrical signals that induce cell death.

## Results

### Starving populations of hyposocial *B. subtilis* are rapidly lysed by monovalent cations

By serendipity we discovered that a mutant strain with impaired chemical communication (relevant genotype: *ΔcomA ΔrapCphrC ΔrapFphrF* P_*hyperspank*_*-*P_*comA*_*-comA*D55A), and for simplicity denoted as “hyposocial” strain, is highly susceptible to lysis in phosphate-buffered saline (PBS) (Fig. 1a). Suspensions of this strain in PBS, an isotonic salt solution that is commonly used for handling bacteria in the laboratory, lost 50% of their optical density (OD_600nm_) within an hour (T_50_ = 1.0 ± 0.1 h), while in Tris-buffer (Tris-HCl, pH 7.4) lysis occurred at a slower rate (T_50_ = 3.0 ± 1h) (Fig. 1b). To identify the stress factor(s) that cause population loss, we determined T_50_ values for cells suspended in Tris in the presence of different salts, osmolytes and nutrients (Supplementary Fig.1). Cell suspensions containing monovalent cations (Na^+^, K^+^, NH_4_^+^) showed a rapid decline in OD, with little difference between the cations tested and regardless of the anion involved (Cl^-^, SO_4_^-2^, PO_4_^-3^, NO_3_^-^); in contrast, the non-ionic osmolyte glycine betaine supplied at isoosmolar concentrations did not induce lysis. Population loss occurred in salt concentrations <150 mM and was decelerated at lower concentrations of monovalent cations. Notably, addition of divalent cations (Mg^2+^, Ca^2+^) prevented lysis, and even halted ongoing lysis by NaCl practically immediately (Fig. 1c). The presence of small amounts of glucose also prevented lysis upon exposure to NaCl (Supplementary Fig. 1). Together, these results imply that, in a nutrient-depleted environment, monovalent cations cause rapid lysis of a hyposocial *B. subtilis* strain.

**Figure 1.**
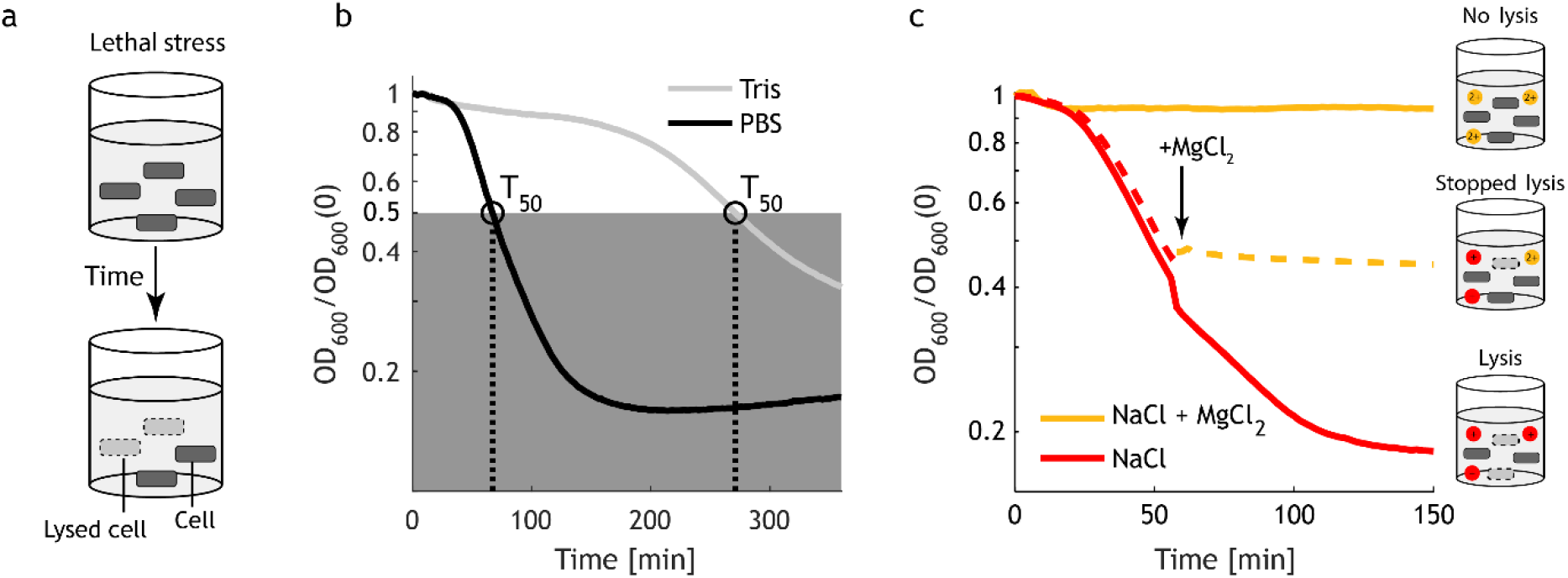
Monovalent cations cause rapid population loss in cultures of a hyposocial *B. subtilis* strain under conditions of nutrient deprivation. **(a)** Scheme illustrating cell lysis of hyposocial *B. subtilis* upon exposure to lethal stress conditions as generated by suspension of cells in phosphate-buffered saline (PBS). **(b)** A hyposocial *B. subtilis* strain lyses quickly. The plot shows the decline in the optical density (OD_600_) as a function of time in PBS (black curve) and in Tris (grey curve). Data was normalized to the initial OD_600_. T_50_ denotes the exposure time required to reduce the initial OD by 50%. Data: mean ± std, *n*_r_ = 6 technical replicates. Strain: BIB1933. **(c)** Monovalent cations induce lysis, while addition of divalent cations prevents (and rapidly stops ongoing) lysis. Representative OD curves for cells suspended in Tris and exposed to NaCl in the absence (red), presence (orange) or upon acute addition of MgCl_2_. Strain: BIB2153.

### ComA signaling mutants have pleiotropic effects on stress behavior and cell morphology

Cell-cell communication was not previously known to be required to protect against ionic stress. The transcription factor ComA occupies a central position in the *B. subtilis* communication network (Fig. 2a). ComA is regulated by several signaling systems – by the pheromone ComX, which affects its phosphorylation status via the histidine kinase ComP^20,21^, and by sequestration by Rap-modulator proteins^22,23^ that are inhibited by signaling via a number of Phr peptides, respectively. Notably, there are no obvious direct or indirect gene targets of ComA that could confer resistance^24,25^. Surprisingly, signaling mutants that deregulate ComA activity showed pronounced differences in their ability to withstand lysis in PBS. A subset of mutant strains was more prone to lysis than the wild type, as evidenced by their lower T_50_ values, while other mutants were even more resistant than wild-type cells (Fig. 2b). In the hyposocial strain lysis occurred twice as fast (T_50_ = 55±5 min) as in the wild type (T_50_ = 101±8 min). In contrast, a mutant strain which responds sensitively to the ComX pheromone due to mutations that favor the accumulation of phosphorylated ComA (relevant genotype: *ΔcomA ΔrapCphrC ΔrapFphrF* P_*hyperspank*_*-*P_*comA-*_*comA*), and for simplicity denoted as “hypersocial” strain, lysis was considerably delayed (T_50_ = 160±30 min). In addition to this difference in stress behavior, these two mutants differed in their morphologies (Fig. 2c). The hyposocial strain primarily contained short rod-shaped cells (that exhibited vigorous movement when immersed in liquid films; not shown), while the hypersocial strain was dominated by chains of connected cells (which were non-motile; not shown) that were reminiscent of sessile cells.

**Figure 2.**
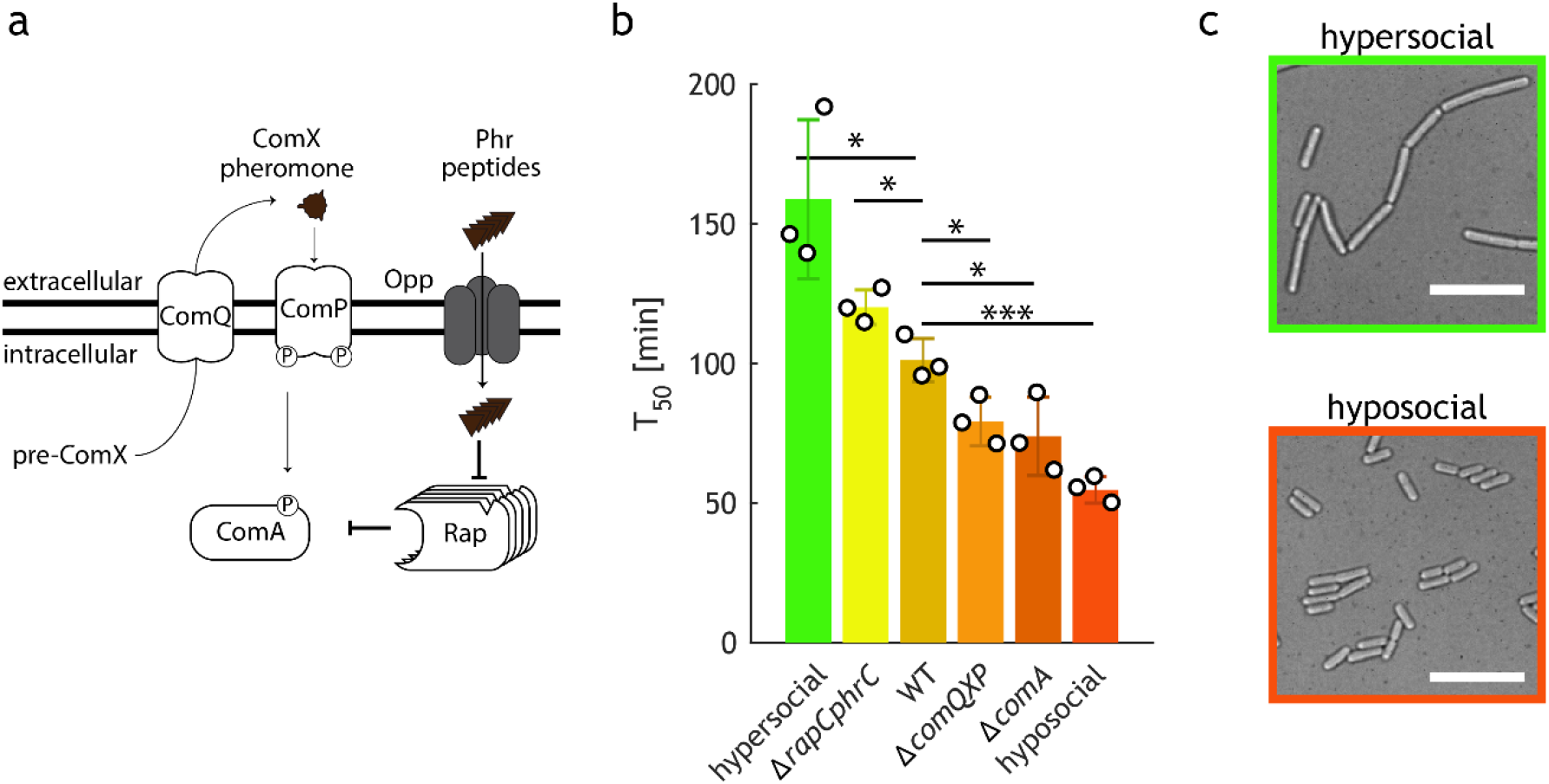
Signaling mutants exhibit pleiotropic effects on stress tolerance and cell morphology. **(a)** Simplified scheme of the ComA regulatory network. Upon binding of the pheromone ComX, the histidine kinase ComP phosphorylates ComA. Rap receptors are controlled by Phr signaling peptides. Raps sequester ComA and this inhibition is relieved upon binding of Phrs to their cognate Rap-receptors. **(b)** T_50_ values for the indicated ComA-regulatory mutant strains in PBS. From left to right BIB1931, BIB351, BIB224, BIB2036, BIB2038, BIB1933. Data: Bars represent means ± SD for *n*_e_ = 3 independent experiments (circles). Unpaired *t* test: \**P*≤0.05, \*\**P*≤0.01, \*\*\**P*≤0.001. **(c)** Bright-field micrographs of cells derived from the hypersocial (top, BIB2109) and hyposocial mutant (bottom, BIB2107) show different cell morphologies. Scale bar: 10 µm.

### Sodium ions kill motile cells

The pleiotropic effects of deregulated ComA activity on stress behavior and cell morphology raised the possibility that the two cell types differ in their resistance traits. To investigate this, we engineered wild-type *B. subtilis* carrying fluorescent reporters that specifically mark motile and sessile cells by fusing the promoter of the *hag* gene (coding for flagellin) to YFP^4^ and the promoter of the *tapA* gene (coding for a protein involved in extracellular matrix synthesis) to CFP^26,6^, and used fluorescence time-lapse microscopy experiments to study their stress responses at the cellular level (Fig. 3a). Wild-type populations comprised a mix of both cell types: short rods expressed YFP from the *hag* promoter (displayed in magenta) while cells in chains expressed CFP from the *tapA* promoter (displayed in cyan) (Fig. 3b, left). For simplicity, we will refer to “*hag* on” cells as “motile” and “*tapA* on” cells as “sessile”, although both cell types are immobilized on Tris-based agarose pads in our experiments. Exposure of cells to NaCl selectively eliminated the motile subpopulation of cells (Fig. 3b, right, Supplementary Movie S1). Motile cells lysed rapidly and their numbers decreased exponentially at a rate of *λ*_*m*_ = (3.5 ± 1.0) h^−1^ (corresponding to a T_50_ = 13±4 min) resulting in the loss of more than 95% of all cells within an hour. In contrast, sessile cells did not lyse and their numbers remained stable over the same time period (Fig. 3c). We next recorded the response of each cell type to exposure to NaCl and subsequent addition of nutrients (Fig. 3d). The vast majority [99(±1)%] of motile cells underwent lysis, and only a negligible fraction survived the stress and resumed growth when nutrients became available. In contrast, the majority [50(±7)%] of sessile cells grew and another 14(±1)% were classified as “inert”, i.e. these cells did not lyse, but they also failed to initiate visible growth within 30 min following nutrient addition. In a sodium-free environment little lysis of either cell type occurred (see also Fig. 3c, Inset) and 82(±5)% of motile and 94(±6)% of sessile cells were able to grow upon nutrient supplementation. Together, these data show that NaCl preferentially kills motile cells.

**Figure 3.**
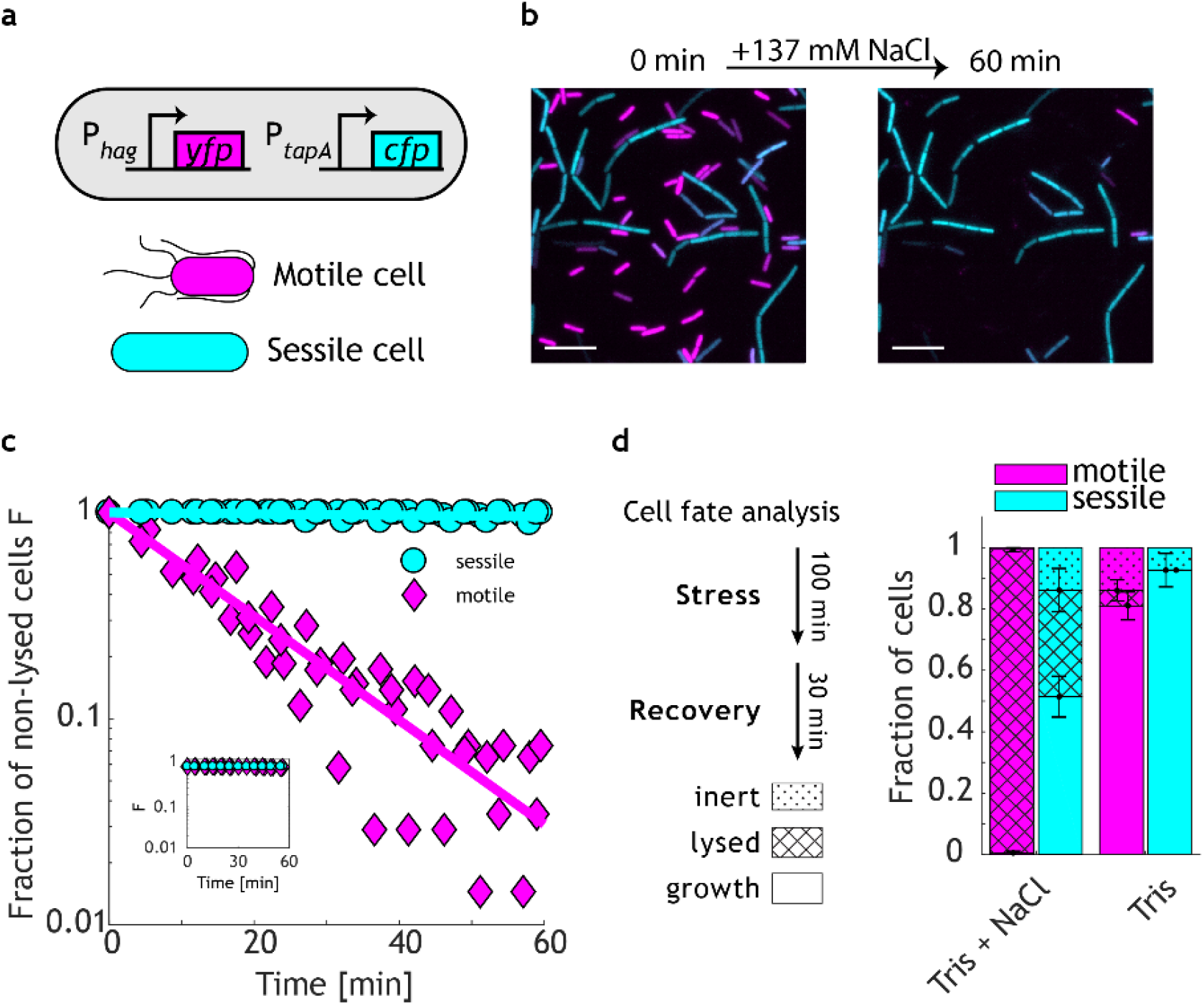
Na^+^ is highly lethal to short, rod-shaped motile cells. **(a)** Scheme of the cellular reporter system used to fluorescently label motile and sessile cells. Motile cells activate the *hag* promoter and express YFP. Sessile cells activate the *tapA* promoter and express CFP. **(b)** Fluorescence micrographs of a population of reporter cells (in an otherwise wild-type background, BIB2127) before and after exposure to NaCl. Left: The initial population is composed of short, rod-shaped cells expressing YFP (false colored in magenta) and chains of cells expressing CFP (colored in cyan). Right: NaCl selectively eliminates the YFP-expressing motile subpopulation. Scale bar: 10 µm. **(c)** Fraction (*F*) of non-lysed cells in motile (magenta) and sessile subpopulations (cyan) plotted as a function of time after exposure to NaCl, as determined by time-lapse microscopy. Data: n = 4 movies with at least N_c_ > 24 cells for each cell type. The magenta line denotes a fit which describes lysis dynamics as an exponential decay (T_50_ = 13±4 min). Inset shows corresponding control without NaCl. **(d)** NaCl kills motile cells. Left: Illustration of the protocol used for the analysis of cell fate. The fate of single cells was recorded after exposure to NaCl for 100 min and subsequent addition of nutrients (LB) for 30 min. Cells were classified as lysed, growing or inert (non-lysed, but non-growing), and the relative proportions of each detected among motile (magenta bars) and sessile subpopulations (cyan bars) with and without exposure to NaCl are shown on the right in a stacked column chart. Bars represent weighted means ± weighted std of *n*_r_ = 4 technical replicates, covering in total at least *N*_c_ = 270 cells of each type.

### Differential expression of *lytABC* is responsible for cell-type specific lysis

The major autolysin LytC is required for cell lysis under a variety of stress conditions^10,11,19,27,28^. LytC is a cell-wall amidase that is encoded in an operon, and expressed together with the enhancer LytB and the lipoprotein LytA. Expression is controlled by two promoters, one of which is dependent on the cell-type-specific sigma factor SigD, which is active in motile cells^29–31^. We therefore asked whether differential expression of *lytABC* contributes to cell-type-specific cation-induced lysis. Indeed, deletion of the *lytABC* operon largely prevented lysis (Fig. 4a). Microscopic investigations using the cell-type-specific markers revealed that the mutant retained both cell types and that lysis of motile mutant cells upon exposure to NaCl was inhibited (Supplementary Fig. 2). We also monitored *lytABC* gene expression with a fluorescent promoter fusion to mCherry in a wild-type background. As expected, the distribution of mCherry fluorescence intensity obtained from motile cells was shifted to higher values relative to those in sessile cells (Fig. 4b). If differential *lytABC* expression was indeed responsible for cell-type-specific lysis, it should be possible to reverse the lysis pattern by directing *lytABC* expression to sessile cells. We therefore engineered the *lytABC* deletion strain to ectopically express the *lytABC* operon (in tandem with mCherry) under the control of the *tapA* promoter. Indeed, in the engineered strain, mCherry fluorescence was higher in sessile cells compared to motile cells, suggesting that the *lytABC* expression pattern was successfully reversed (Fig. 4c). Moreover, lysis was now preferentially observed in sessile cells, as indicated by the dissolution of cell walls (Supplementary Movie S2) and a sharp drop in cellular fluorescence in CFP-expressing sessile cells (Fig. 4d).

**Figure 4.**
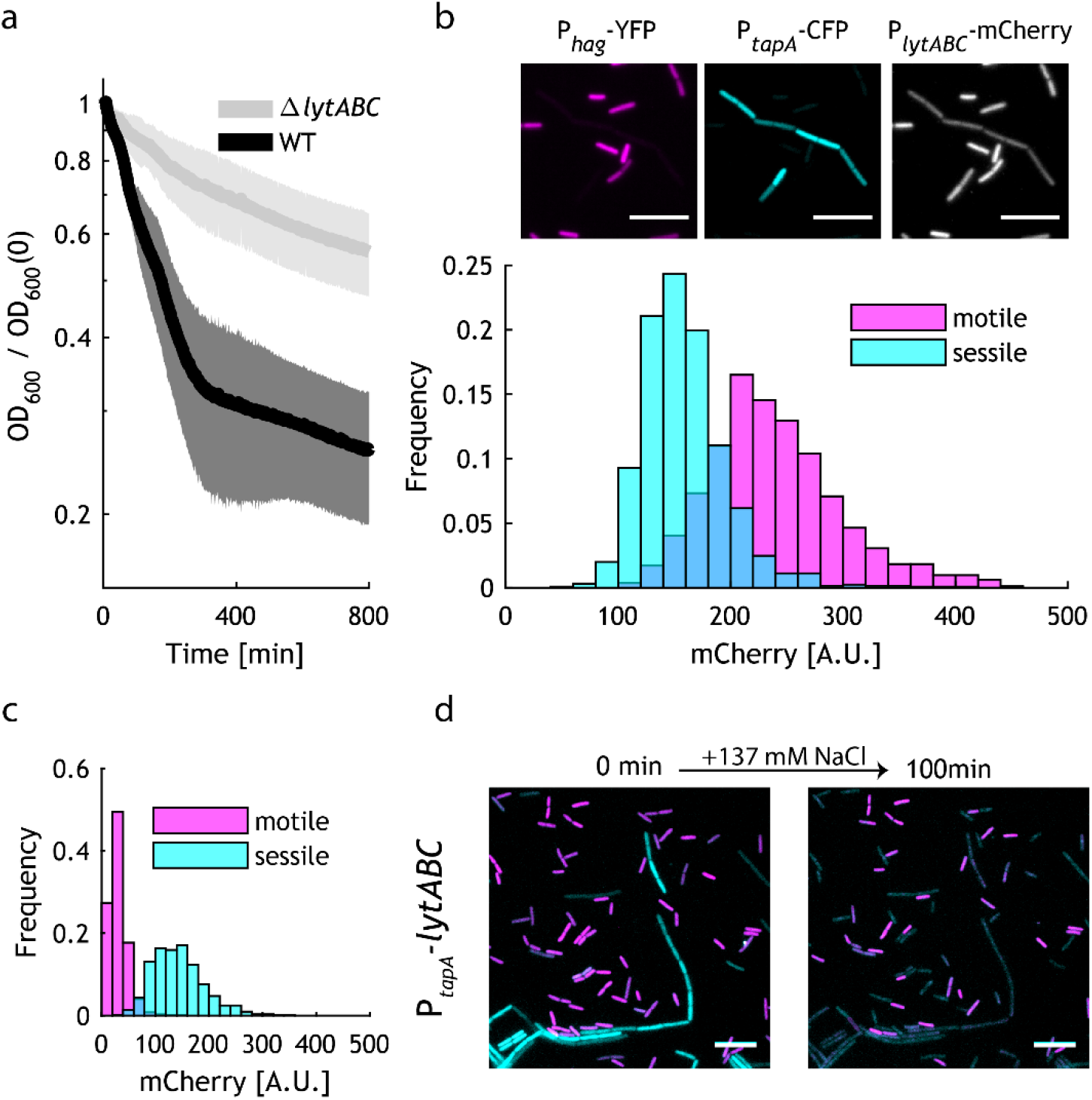
Differential expression of *lytABC* results in cell-type-specific lysis. **(a)** Deletion of *lytABC* prevents population loss in Tris+NaCl. Data: mean ± std of *n*_e_ = 3 independent experiments. Strains: BIB1937, BIB224. **(b)** Motile wild-type cells show elevated levels of expression from P_*lytABC*_ compared to sessile cells. Top: Fluorescence micrographs of wild-type cells carrying a P_*hag*_-YFP P_*tapA*_-CFP P_*lytC*_-mCherry triple reporter (BIB2246). From left to right: YFP, CFP and mCherry channels. Scale bar: 10 µm. Bottom: Corresponding histograms of cellular mCherry fluorescence intensities for motile (magenta, *N*_c_= 818) and sessile (cyan, *N*_c_= 1249) subpopulations obtained from 20 images. **(c)** The cell-type-specific expression pattern of *lytABC* can be reversed. Engineered sessile cells (BIB2356) carrying a dual P_*hag*_-YFP P_*tapA*_-CFP reporter express *lytABC* together with mCherry. Histograms of cellular mCherry fluorescence intensities for motile (magenta, N_c_ = 621) and sessile (cyan, *N*_c_ = 527) subpopulations obtained from *n*=3 images. **(d)** NaCl lyses reprogrammed sessile cells. Micrographs of BIB2356 before (left) and after exposure to NaCl (right). Images show superposition of the YFP (magenta) and CFP (cyan) signals. Scale bar: 10 µm.

### LytABC sensitizes cells to stress and mediates cation-induced cell death

We next asked whether the action of LytABC is restricted to the elimination of non-viable cells, or whether it might also sensitize cells to stress and thereby mediate stress-induced cell death. We also wondered whether sessile cells have additional layers of protection or could be similarly sensitized to stress by raising LytABC levels. To address these questions, we engineered an IPTG-inducible *lytABC* operon into the *lytABC* deletion strain carrying the cell-type specific markers and tracked the fate of each cell upon exposure to NaCl stress and the subsequent addition of nutrients (Fig. 5a). Without induction, i.e. when cells had only leaky expression of LytABC, the majority of motile cells (55±16%) survived the stress and resumed growth albeit at slightly lower frequencies than sessile cells (67±20%). All remaining cells were inert and did not lyse (Supplementary Movie S3). In contrast, upon full induction of *lytABC* by IPTG, both motile (83±7%) and sessile cells (72±3%) lysed (Supplementary Movie S4). The percentage of growing cells dropped to 17±7% in motile and 28±3% in sessile cells, respectively. Moreover, in both cell types, the fraction of lysed cells observed following full induction of *lytABC* expression exceeded the fraction of inert cells under uninduced conditions by roughly a factor of two. We thus conclude that LytABC sensitizes cells to stress and contributes to cell death in motile and, when overexpressed, also in sessile cells.

**Figure 5.**
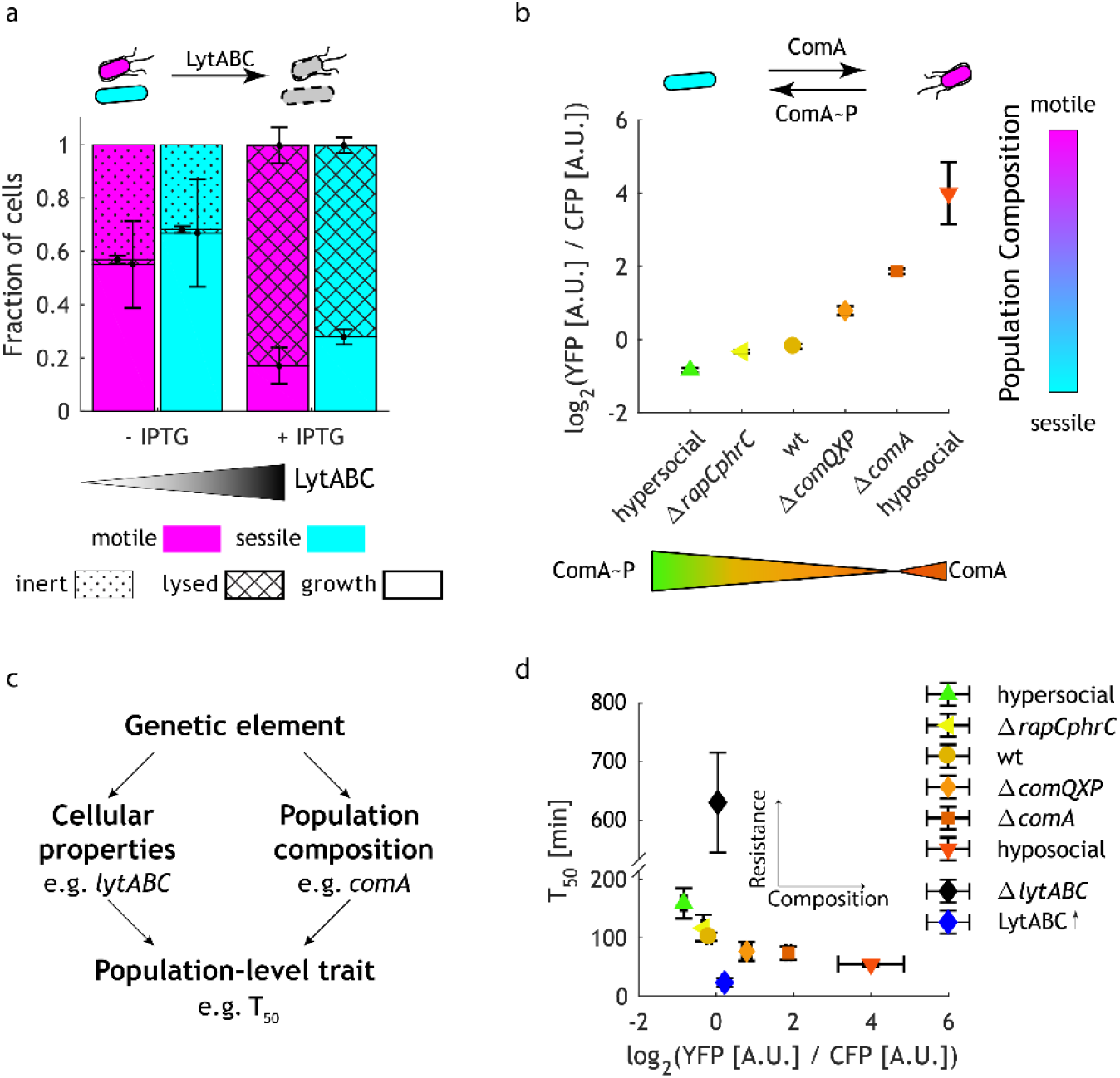
*lytABC* and *comA* control fitness at the cellular and population levels, respectively. **(a)** LytABC enhances susceptibility to monovalent cation stress in both cell types. The stacked column chart summarizes the results of a cell-fate analysis (after 60-min exposure to NaCl, followed by 60 min recovery in LB*) for motile and sessile subpopulations that were engineered to express *lytABC* upon IPTG induction. Strain: BIB2330. Data: weighted mean ± std, n_r_ = 3 replicates. **(b)** ComA controls population composition. Results of population composition measurements based on cell-type-specific marker constructs in the indicated mutant strains using the r=log_2_(YFP/CFP) of the bulk fluorescence as a proxy. Mutations that increase ComA∼P levels shift the population towards the sessile state. Mutations that increase ComA levels shift the population towards the motile state. Data: mean ± std, n_r_ = 8 replicates. Strains: BIB2107, BIB2129, BIB2127, BIB2135, BIB2137, BIB2109. **(c)** Principal functions of the genetic elements involved in controlling population-level traits (such as stress-induced population loss as quantified by T_50_ values). A genetic element can either directly alter a cell’s resistance to stress, or alter the composition of a population comprised of two cell types with different resistance properties (or both). **(d)** Effects of the indicated mutations on T_50_ values in PBS. Genetic perturbations of social signaling modulate fitness by altering the composition of the population, while *lyt* mutants shift T_50_ without affecting composition. T_50_ replotted from Fig. 2C. log_2_(YFP/CFP) replotted from Fig. 5B. Δ*lytABC* P_hyperspank_ *lytABC* (BIB2330) without induction; LytABC↑: BIB2330 with 1mM IPTG.

### ComA has a dual function in modulating population composition

To assess how ComA affects population composition, we transformed all mutant strains with the P_*hag*_-YFP P_*tapA*_-CFP cell-type reporter. Fluorescence microscopy experiments suggested that fluorescence from both promoters provides a relatively stable marker for cell type, regardless of genotype (Supplementary Fig. 3). Since measurements of subpopulation composition by microscopy turned out to be unreliable (due to the patchy distribution of both subpopulations on agarose pads), we used the population-averaged ratio of fluorescence as measured in a fluorescence plate reader as a proxy for population composition, *r*=log_2_(YFP/CFP). *r* varied with genotype (Fig. 5b). All genetic perturbations that were expected to increase the levels of ComA∼P decreased *r*, suggesting that ComA∼P favors the sessile state. In contrast, in a *comA* deletion strain, *r* was shifted to higher values than the wild type, indicating that the population had a higher proportion of motile cells. Surprisingly, the hyposocial mutant, which expresses non-phosphorylatable ComA due to an alanine substitution at its phosphorylation site D55 and which lacks RapF- and RapC-dependent ComA sequestration, showed an even higher *r* than the *comA* deletion mutant, suggesting that accumulation of unphosphorylated ComA favors the motile state.

### *lytABC* and *comA* control fitness by regulating stress tolerance and population composition, respectively

Taken together, our data show that both *lytABC* and *comA* (and related genes) have a profound effect on fitness under stress, but they act in very different ways, namely by altering cellular properties and population composition, respectively. Quantitative characterization of population-level traits in bacteria is in its infancy, and their underlying genetic bases have yet to be understood. In principle, any gene could affect a population-level trait in two ways: either by changing the behavior of individual cells or by adjusting population composition without altering the individual properties of cells or both (Fig. 5c). To distinguish between these possibilities, we analyzed effects on T_50_ in conjunction with the quantification of population composition by measuring log_2_(YFP/CFP) ratios in reporter populations (Fig. 5d). As expected, increased expression of LytABC decreased T_50_ values without significantly affecting population composition. In contrast, for *comA* and related genetic perturbations to social signaling the inferred changes in population composition with altered genotypes were highly correlated with the T_50_ values that were obtained from population-level lysis experiments: the higher the fraction of sessile cells in the population, the more resilient the population. Moreover, a mathematical model that assumes that these genetic mutations alter population composition without affecting the lysis properties of each cell type described the population dynamics in the complete mutant dataset very well (Supplementary Fig. S4). The model inferred a roughly ten-fold difference in the lysis rates for the two subpopulations (*λ*_m_ = 1.5 h^−1^ λ_s_ = 0.1 h^−1^), which is in accordance with the single-cell experiments that show that sessile cells are much more tolerant to stress than motile cells. We thus conclude that hypersocial strains have a fitness advantage under stress due to an increased fraction of sessile cells.

## Discussion

The generation of phenotypic population-level heterogeneity is an important strategy that bacteria employ as part of their adaptive repertoire to survive in nature. *B. subtilis* is known to diversify phenotypically under a variety of conditions, including planktonic cultures^4^ and biofilms^8^. However, this factor has been - and often still is - neglected when analyzing resistance traits. A variety of stress conditions are known to result in cell lysis^10–19^. Notably, some studies have reported the “normal” cell shape of *B. subtilis* as “short rods”^32^ while others observed cells growing in chains as being dominant in their experimental setting^15^. In many other cases, populations appeared to be comprised a mixture of both cell types, but this elicited no comment^10,18^. Our work clearly shows that motile and sessile cells have different resistance properties, and population-level resistance traits will therefore depend on population composition. Strain background, cultivation conditions and growth phase can affect population composition even in exponentially growing populations, making it difficult to compare different studies. We therefore suggest that all future studies should carefully characterize the composition and/or cell state of their populations.

The transition between motile and sessile cell states is regulated by a complex regulatory network centered around the SinR switch^1,2^, which is deeply embedded within the *B. subtilis* regulatory network. Thus, many genes with documented pleiotropic effects are expected to affect population-level traits, changing population composition rather than changing cellular properties. For example, the lysis-resistant isolate FJ2 exhibits cellular characteristics of bacteria locked in the sessile state, and indeed a mutation was mapped to *sinR*. The mutant strain did not lyse when exposed to surfactant stress^12^, cold shock^19^ or ion stress^17^. On the other hand, the seminal work by Joliffe et al.^14^ on cellular autolysis was performed on a *spo* mutant strain that was probably unable to generate sessile cells – which presumably contributed to its invariably low stress tolerance. It will undoubtedly be a challenging but vital task to rigorously and comprehensively measure effects on cellular traits, as well as population composition on a genome-wide scale in order to clearly assign gene functions.

### ComA-dependent cell-type switching may confer benefits that stabilize cooperative traits

Quorum sensing is a mode of orchestrating population-level behavior, which is commonly used in the biosphere to regulate the production of public goods. Interestingly, a growing body of evidence indicates that quorum sensing is tightly linked to stress tolerance. In several Gram-negative bacteria, quorum-sensing-deficient mutants have been shown to be vulnerable to various stress factors, including osmotic^33,34^, oxidative^35^, thermal^33^ and heavy-metal stress^33^ and others. In *Vibrio harveyi*, increased stress tolerance results from direct regulation of stress-response pathways by a regulator that is controlled by quorum sensing^34^. Coupling of quorum sensing to stress resilience confers a private benefit to quorum-sensing-proficient cells, and thereby can contribute to stabilization of social investments and help to eliminate cheaters^33^. Similar reasoning should also apply to *B. subtilis*. ComA directly controls the production of two public goods (surfactin and pectate lyase); however, a direct regulation of stress-response pathways by ComA seems unlikely^24,25^. Nevertheless, our data suggests that mutants with impaired pheromone-signaling are less fit under cation stress due to an altered population composition. Together with the observation that similarly impaired strains also fail to differentiate into competent (or K-state) cells^36^and endospores^37^respectively, this implies that an inability to communicate leads to a fitness disadvantage under a variety of stress conditions.

Our mutant data clearly show that ComA controls the composition of vegetative *B. subtilis* populations, and therefore must modulate switching between the motile and the sessile cell state. Phosphorylated ComA∼P and non-phosphorylated ComA seem to have opposing functions and favor the sessile and motile states, respectively. How this is achieved at the molecular level requires further study. The promoters of ComA target genes have surprisingly complex architectures^24^, which might facilitate differential activation by ComA and ComA∼P; and at least some target genes have (indirect, higher-order) connections to SinR, the regulator of the lifestyle switch^6,25^. Since deletion of *comQXP* reduced the frequency of sessile cells in our populations, ComX-based signaling seems to favor transitions to the sessile state. However, we note that additional way(s) to favor the sessile state must exist beyond the previously proposed activation of surfactin synthesis via ComA∼P^6^, since the B168 laboratory strain that was used in our study is deficient in surfactin synthesis^38^. Previous (population average) transcriptome data for *comA* mutants also support the notion of a ComX-dependent, but surfactin-independent transition to the sessile state, as transcripts of genes coding for matrix production were underrepresented^25^.

### Elevated autolysin expression in motile cells may reflect a tradeoff

Our data show that sessile cells are more tolerant to stress but, perhaps surprisingly, these cells were not necessarily better protected from stress; instead, motile cells were more vulnerable as a result of elevated expression of the *lytABC* operon. When the operon was artificially induced in sessile cells they became (almost) as vulnerable to the same stresses as motile cells, while deletion of the *lyt* operon enhanced the resistance properties of motile cells to the levels seen in sessile cells. Motile cells activate *lytABC* from a cell-type specific SigD-dependent promoter to raise the levels of the autolysin complex. This regulatory feature should therefore confer a direct benefit on motile cells. Indeed, although *lytABC* is not required for swimming *per se*^11^, *lytC* is required for proper flagellar function and swarming^11,3^. Moreover, LytC, together with other components of the motility apparatus, contributes to the formation of nanotubes, which can connect cells together, and facilitate material transfer between them^39^.

How LytABC contributes to these functions is not completely understood. The *lytABC* operon codes for a cell-wall lytic complex comprised of the LytC amidase, the enhancer LytB and the lipoprotein LytA. *In vitro* the rate of cell-wall hydrolysis by LytC increases in the presence of monovalent cations, is reduced by divalent cations and is also affected by pH^17,40^, suggesting that ions alter enzyme function and/or cell-wall organization, and thereby modulate enzyme-substrate interactions. In living cells, the distribution of ions and protons in the cell wall is affected by both external and internal factors. Specifically, growing *B. subtilis* cells generate a protonmotive force (PMF), which results in a protonated cell wall^41^. In motile cells the PMF is locally dissipated to drive the flagellar motor, which may increase the demand for cell-wall remodeling. Regardless of the underlying mechanism, the elevated LytABC levels in motile cells probably reflect a tradeoff that ensures that the complex can carry out its beneficial functions in motile cells under relaxed environmental conditions, albeit at the price of a reduced stress tolerance.

Many stress conditions, including entry into stationary phase^27^, biosurfactants^28^, cold shock^19^, sodium azide^11^ and oxygen depletion^10^ are known to result in LytC-dependent autolysis. Importantly, as we have shown here, LytABC not only eliminates non-viable cells, but also contributes to the induction of cell death under starvation conditions. Recent data reported by Arjes et al. shows that a *lytC* mutant strain has a somewhat increased viability relative to the wild type when exposed to oxygen depletion^10^. Moreover, upon closer inspection of their video material, the wild-type population seems to be made up of a mixture of motile and sessile cells. Surviving cells predominantly grow in chains, and therefore probably represent mostly sessile cells. Others have also suggested that autolysins might contribute to killing, based on a higher viability of the lysis-resistant mutant strain FJ2 to exposure to surfactants^12^. Clearly, the function of LytABC in mediating cell death needs further study and additional mechanisms that reduce viability in response to stress conditions must exist. Yet it seems likely that motile cells are more susceptible than sessile cells to a variety of stress factors, specifically when these are encountered in conjunction with monovalent cations.

### Electrical signaling may trigger the death of motile cells

The vulnerability of motile cells to monovalent cations might be profitably exploited by sessile cells. Monovalent cations (specifically potassium ions) have recently been proposed to facilitate “electrical communication” within *B. subtilis* biofilms, in which potassium waves are elicited by metabolically stressed cells in the interior to restrict biofilm expansion at the periphery^42,43^. Moreover, the electrical activity of biofilms correlates with an accumulation of motile cells (*B. subtilis* and other species) at the edge of the biofilm^44^. The ion concentrations that were used in these experiments are comparable to those employed here, raising the possibility that the cation signals not only trap motile cells, but could also help to kill them. Our data clearly show that LytABC facilitates killing by sodium ions when motile cells are starved. We also observed cell-type-selective killing with potassium ions and in an undomesticated *B. subtilis* strain (Supplementary Movies S5 & S6). In order for monovalent cations to kill, they probably have to act in concert with stress factors, such as starvation^14^, oxygen depletion^10^, toxins^15^ or surfactants^12^, which dissipate the protonmotive force and/or disturb the membrane potential, respectively. If so, motile cells (of *B. subtilis* and likely also other species) could become prey for hungry *B. subtilis* sessile cells that elicit electrical signals to kill, lyse and feed on these cells^15,45,46^. In addition to that, *hag*-expressing and thus presumably motile *B. subtilis* cells are present at distinct locations within biofilms^8^. It is thus conceivable that cation-induced lysis of this subpopulation could contribute to the specific cell-death-based patterning that drives biofilm “morphogenesis” by projection of aerial structures^47^.

### Applications

From a practical point of view, the exquisite sensitivity of subpopulations of bacteria to salt buffers such as PBS raises a strong caveat with respect to its use in experimental protocols. Moreover, in order to improve stress tolerance of strains used in biotechnological applications *lytABC* is an obvious candidate for deactivation. On the other hand, cation-sensitive lysis by LytABC could be useful for some synthetic biology applications: targeting *lytABC* expression may allow for precise elimination of any – even a transient – cell state, as demonstrated here. Finally, other both Gram-positive and Gram-negative bacteria have been reported to undergo autolysis in conditions similar to ours^48–51^, raising the possibility that proteins that function analogous to LytABC are widespread.

## Materials and Methods

### Strains

All strains were derivatives of the laboratory strain 1A700 (W168) and their genotypes are listed in Supplementary Table S1. Experiments were also performed with PS216, a recent natural isolate^52^.

### Plasmid construction

All plasmids and oligonucleotides used in this study are listed in Supplementary Tables S2 and S3, respectively. *E. coli* DH5α was used for cloning. Plasmids were constructed by ligation-independent cloning (LIC) and by restriction-enzyme ligation cloning (RELC), respectively. All DNA fragments were amplified from the genomic DNA of *B. subtilis* W168 (unless noted otherwise). Plasmids were verified by analytical restriction digestion and sequencing.

Plasmids for fluorescent promoter fusions were constructed by amplifying 100-300 bp of the promoter region located upstream of the ribosome-binding site, and inserted into pXFP_Star and pXFP_bglStar, respectively. pXFP_bglStar vectors were constructed by RELC from pXFP_Star^53^ and the backbone of pXFPbglS^54^ using EcoRI digestion and have a TgyrA terminator included upstream of the LIC site for promoter insertion to buffer spurious transcription. The new pXFP_bglStar vectors have been deposited in the Bacillus Genetic Stock Center (Columbus, OH, USA) under the Accession Nos. ECE793 (YFP), ECE794 (GFP) and ECE795 (CFP).

Plasmids for gene deletions were constructed by amplifying 500-bp – 1-kb fragments located upstream and downstream of the region of interest. Fragments were fused by PCR and cloned into pMAD^55^ by RELC.

Plasmids for inducible gene expression were constructed by amplifying genes and operons of interest and inserted into pDR111 (David Rudner, Harvard University, Boston) and pDG1662^56^, respectively, by RELC. In order to mimic native expression of ComA under uninduced conditions, P_*comA*_-*comA* was cloned into pDG1662. The expression vector EIB517 for ComAD55A was obtained by site-directed mutagenesis of EIB281.

To construct the plasmid used for differential gene expression of the *lytABC* operon in sessile cells (EIB799), the P*tapA* promoter (amplified from EIB727) was fused to an optimized ribosome-binding site and the *lytABC* operon by PCR, and the resulting construct was inserted into pRFP_Star by LIC.

### Strain construction

Strains were constructed by a one-step^57^ or a two-step transformation method^58^ with the ectopic integration vectors listed in Supplementary Table S3. The pLK-comK plasmid was used to transform strains with reduced competence development and the final strains were cured from the plasmid using standard protocols^59^. All reporters and expression cassettes were integrated at an ectopic locus (*amyE, bglS*) and present in single copy as verified by PCR. We also checked that transformations carried out with pDR111 retained an intact *ldh* locus^60^. Clean gene deletions were constructed using pMAD plasmids, following a protocol similar to that of Arnaud et al.^55^. At each step, we verified by appropriate PCRs that the gene had been deleted from the chromosomal locus, that the pMAD plasmid had been lost and, finally, that the gene was absent from the chromosome.

### Media and solutions

*B. subtilis* strains were grown in LB media (10g l^-1^ Tryptone, 5g l^-1^ yeast extract, 5g l^-1^ NaCl) and S7_50_ medium^61,62^. The appropriate antibiotics were added as required: chloramphenicol (5 µg/ml), kanamycin (10 µg/ml), spectinomycin (100 µg/ml), or a combination of erythromycin (1 mg/ml) with lincomycin (25 µg/ml); X-Gal was used at 100 µg/ml, and IPTG (isopropyl-β-D-thiogalactopyranoside) at 100 µM.

Solutions for stress experiments were performed with autoclaved phosphate-buffered saline (PBS) (137 mM NaCl, 2.7 mM KCl, 10 mM Na_2_HPO_4_, 2 mM KH_2_PO_4_, pH=7.4) and filter-sterilized Tris (10 mM, adjusted to pH 7.4 with HCl) or Tris-based solutions (10 mM Tris, pH 7.4, added at the indicated concentration). Where indicated, supplements were added during preparation and pH was adjusted with HCl. Divalent cations were added from 2M aqueous solutions of MgCl_2_ and CaCl_2_, respectively.

Microscopic analyses of killing and cell fate were performed with aqueous stock solutions of 2.2 M NaCl or 2.2 M KCl to induce cation stress, and 4x concentrated LB medium lacking NaCl to facilitate cell recovery.

### Cell culture

*B. subtilis* strains were inoculated from single colonies on LB plates into 5 ml LB with antibiotics, and grown in culture tubes (18 mm) at 37°C and 180 rpm in a Infors Multitron II for 6 h at 37°C. Cells were resuspended in 3 ml fresh S7_50_ medium with antibiotics to a starting optical density at 600 nm of 0.02, and grown overnight (18 h) in culture tubes (16 mm) at 30°C. Samples were diluted into 10 ml of fresh S7_50_ medium (OD = 0.02) without antibiotics, and grown in 100 ml flasks at 180 rpm for 6 h at 37°C in an Infors Ecotron.

Cells were harvested by centrifuging at 3273 rcf for 5 min (for plate-reader experiments) and at 16-17 rcf for 1 min (for microscopy experiments) and washed once. For initial population-loss experiments involving PBS, cells were washed in PBS. For all other experiments cells were washed in Tris-HCl to reduce the likelihood of cell lysis during washing. Cells were resuspended in Tris-HCl to OD = 2.0 (for microscopy experiments) and to OD = 2.5 in the indicated solution (for plate-reader experiments).

### Plate Reader Assays

Population loss was monitored by absorbance measurements in 96-well plates (96 MicroWell™, Nunc™, Thermo Scientific) using at least n = 6 technical replicates with 200-µl sample volumes. Plates were incubated at 37°C in a preheated Epoch 2 Microplate Spectrophotometer (BioTek) under linear shaking (1 mm) at 1096 cycles per minute. The OD at 600 nm was recorded at 2-min intervals. Where applicable, the incubation was interrupted to add 10 µl of an aqueous solution of divalent cations to selected wells. Curves were normalized with respect to their initial OD. T_50_ was obtained by linear interpolation between t_i_ and t_i-1_, where t_i_ is defined as the time-point at which normalized OD dropped below 0.5.

Bulk fluorescence was measured in 96-well plates (CellView Microplate, Greiner) using at least n = 6 technical replicates with 150-µl sample volume. Bottom fluorescence was recorded using a fluorescence plate-reader assay (Tecan i-Control Inifinite M1000Pro) at the following settings: YFP (513 / 10 nm, 559 / 10 nm) and CFP (438 /10 nm, 470 /10 nm) at 25 flashes (400 Hz) and 20 µs integration time. Fluorescence was normalized to OD and the OD-normalized background fluorescence from a wild-type strain without reporter was subtracted. Corrected fluorescence was used to calculate log2-ratio r = log_2_ (YFP/CFP).

### Microscopy

Samples (3 μl) of cell suspension (OD = 2.0) were spotted on agarose pads (Tris-HCl solidified with 1.2% Ultrapure Agarose, dimensions: 0.9 cm diameter, 1 mm height) and stamped into 24-well SensoPlates (Greiner). Positions were manually selected to track the fate of both cell types in the same field of view (note that cell types were unevenly distributed on the same pad). Fluorescence and bright-field images were recorded before the onset of stress. To induce cation stress, the plate was removed from the microscope and 4 µl of stress solution was added to the upper surface of the pad and imaging was resumed at the same positions. If applicable, time-lapse movies were recorded by bright-field imaging every 5 min. For cell-fate analysis at least one bright-field image was taken before adding the recovery medium (3 µl) and at the end of the indicated time period.

Imaging was performed on a DeltaVision Elite Imaging System (Applied Precision, Issaquah, WA, USA) equipped with an inverted wide-field fluorescence microscope and encased in an environmental chamber kept at 36.5°C, the Ultimate Focus System and an automated microtiter stage. Cells were imaged with a UPlanSApo 100x/1.40 NA oil objective (Olympus, Tokyo, Japan). Bright-field images were taken at 32% excitation for 0.05 s. Fluorescent reporters were excited by a solid-state illumination unit with a UV filter using the following settings and filters (excitation, emission) and a CFP/YFP/mCh dichroic (reflection bands: 400 nm – 454 nm, 496 nm – 528 nm, 558 nm – 594 nm; transmission bands: 463nm – 487 nm, 537 nm – 550 nm, 602 nm – 648 nm): YFP (513 / 17 nm, 559 / 38 nm), CFP (438 /24 nm, 470 /24 nm) and where applicable, mCherry (575 /25 nm, 632/60 nm) at 100% excitation for 0.5 s each. Images were recorded with a sCMOS PCO Edge camera with 2 × 2 binning using 512 by 512 (Fig. 3, 4b, 4c) or 1024-by-1024 pixels, otherwise with Resolve3D SoftWorx-Acquire Version 7.0.0 Release RC6.

### Image analysis

Fluorescence quantification and cell-type classification: Bright-field images were segmented using a customized software and results were manually inspected. For each cell, the mean fluorescence intensity was determined from the segmented area and the background fluorescence was subtracted^24^. Cells were classified as “motile” (“sessile”) based on fluorescence thresholding. Cells that could not be unambiguously assigned (most likely as a result of a cell-type switching event) were excluded from further analysis.

Cell Fate Analysis: Cells were manually classified as “lysed” if the cell was no longer visible in the bright-field image (or visibly compromised by a dissolving cell periphery), as “growing” if cell length increased by at least 0.5 µm and as “inert” otherwise. Cells that divided during the stress period were excluded from the analysis, unless both daughter cells had the same fate.

## Acknowledgements

We thank Diana Wolf, Stephanie Trauth and Maren Nattermann for construction of *comA* mutant strains and Begonia Ugarte Uribe for construction of the *lytABC* deletion plasmid. We also acknowledge first reports of cell lysis on PBS pads by S. Trauth. We thank M. Knop for providing access to a fluorescence plate reader. We thank Erhard Bremer and Tamara Hoffmann and all members of the Bischofs lab for discussions.

## Author contributions

BS and QC contributed equally to this study. Designed research: BS, QC, IBB. Strain construction: QC, TS, BS. Performed experiments: QC, BS, TS. Analyzed data: BS, QC, IBB. Wrote the manuscript: IBB, BS, QC.

## Funding

This work was supported by the China Scholarship Council (QC), the Landesgraduiertenstiftung Baden-Württemberg in the context of the MMQB program at the BioQuant and in part by the European Research Council (ERC-StG GA 260860) and the Max Planck Society.

## Additional information

**Supplementary information** accompanies this manuscript, including Supplementary Tables 1-3, Supplementary Methods, Supplementary Figure S1-S4 and Supplementary Movies S1-S5 (https://doi.org/10.5446/51550).

## Competing interests

The authors declare no competing financial interests.

## Notes

### Competing Interest Statement

The authors have declared no competing interest.

