## Supplementary Material for "Communication determines population-level fitness under cation stress by modulating the ratio of motile to sessile *B. subtilis* cells"

### SUPPLEMENTARY TABLES

##### Supplementary Table S1. Strains used in this work.

| Strains | Genotype | Source |
|---|---|---|
| ***E. coli*** | | |
| **DH5α** | *$F^{-}$ φ80lacZΔM15 Δ(lacZYAargF)U169 recA1 endA1 hsdR17($r_{K}^{-}$, $m_{K}^{+}$) phoA supE44 thi1 gyrA96 relA1 $\lambda^{-}$* | Invitrogen |
| ***B. subtilis*** | | |
| **BIB224** | *trpC2* | Laboratory stock<br>Orginally obtained from Oscar Kuipers (Groningen) |
| **BIB351** | *trpC2 ΔrapCΔphrC* | This study |
| **BIB398** | *trpC2 pIB69 (pLK-Pspac-comK) replikativ ΔcomA* | This study |
| **BIB1917** | *trpC2 pIB69 (pLK-Pspac-comK) replikativ ΔcomA ΔrapCphrC ΔrapFphrF* | This study |
| **BIB1931** | *trpC2 ΔcomA ΔrapCphrC ΔrapFphrF*<br>*amyE::*[P*hyperspank*-P*comA*-*comA lacI cat*]<br>*bglS::*[TgyrA P*srfAA reshuffled* PIR PDR-*iyfp nptIII*] | This study |
| **BIB1933** | *trpC2 ΔcomA ΔrapCphrC ΔrapFphrF*<br>*amyE::*[P*hyperspank*-P*comA*-*comA*D55A *lacI cat* ]<br>*bglS::*[TgyrA P*srfAA reshuffled* PIR PDR-*iyfp nptIII*] | This study |
| **BIB1937** | *trpC2 ΔlytRABC* | This study |
| **BIB2032** | *trpC2 ΔcomP* | This study |
| **BIB2034** | *trpC2 ΔcomQX* | This study |
| **BIB2036** | *trpC2 ΔcomQXP* | This study |
| **BIB2038** | *trpC2 ΔcomA* | This study |
| **BIB2107** | *trpC2 ΔcomA ΔrapCphrC ΔrapFphrF*<br>*amyE::*[P*hyperspank*-P*comA*-*comA* lacI *cat*]<br>*bglS::*[TgyrA P*tapA*-*cfp* TgyrA P*hag*-*iyfp nptIII*] | This study |

| | | |
|---|---|---|
| **BIB2109** | *trpC2 ΔcomA ΔrapCphrC ΔrapFphrF*<br>*amyE::*[P*hyperspank*-P*comA-comA*D55A *lacI cat*]<br>*bglS::*[TgyrA P*tapA-cfp* TgyrA P*hag-iyfp nptIII*] | This study |
| **BIB2127** | *trpC2 bglS::*[TgyrA P*tapA-cfp* TgyrA P*hag-iyfp nptIII*] | This study |
| **BIB2129** | *trpC2 ΔrapCΔphrC*<br>*bglS::*[TgyrA P*tapA-cfp* TgyrA P*hag-iyfp nptIII*] | This study |
| **BIB2135** | *trpC2 ΔcomQXP*<br>*bglS::*[TgyrA P*tapA-cfp* TgyrA P*hag-iyfp nptIII*] | This study |
| **BIB2137** | *trpC2 ΔcomA*<br>*bglS::*[TgyrA P*tapA-cfp* TgyrA P*hag-iyfp nptIII*] | This study |
| **BIB2153** | *trpC2 ΔcomA ΔrapCphrC ΔrapFphrF*<br>*amyE::*[P*hyperspank*-P*comA-comA* D55A *lacI cat*] | This study |
| **BIB2232** | *trpC2 ΔlytRABC*<br>*bglS::*[TgyrA P*tapA-cfp* TgyrA P*hag-iyfp nptIII*] | This study |
| **BIB2330** | *trpC2 ΔlytRABC*<br>*bglS::*[TgyrA P*tapA-cfp* TgyrA P*hag-iyfp nptIII*]<br>*amyE*::[ *Phyperspank-lytABC lacI spec*] | This study |
| **BIB2332** | *trpC2 ΔlytRABC*<br>*bglS::*[TgyrA P*tapA-cfp* TgyrA P*hag-iyfp nptIII*]<br>*amyE*::[ *Phyperspank-lytC lacI spec*] | This study |
| **BIB2246** | *trpC2*<br>*bglS::*[TgyrA P*tapA-cfp* TgyrA P*hag-iyfp nptIII*]<br>*amyE*::[*PlytABC-mcherry cat*] | This study |
| **BIB2356** | *trpC2 ΔlytRABC*<br>*bglS::*[TgyrA P*tapA-cfp* TgyrA P*hag-iyfp nptIII*]<br>*amyE*::[*PtapA-lytABC-mCherry*] | This study |

**Supplementary Table S2. Plasmids used in this work.**

| Accession Number | Vector | Description | Primers used for cloing | Source |
|---|---|---|---|---|
| **pMAD** | | pBR322-ori pE194-Ts-ori P*clpB*, *bgaB bla erm* | | Ref.[1] |
| **pYFP_bglStar** | | *bglS'* TgyrA LIC promoterless *iyfp kan 'bglS* ColE1 *bla* f1(+) | | This study |
| **pCFP_bglStar** | | *bglS'* TgyrA LIC promoterless cfp$_{bs}$ *kan 'bglS* ColE1 *bla* f1(+) | | This study |
| **pCFPstar** | | *amyE' cat* TgyrA LIC promoterless *cfp*$_{bs}$ *amyE' bla* ColE1 origin | | Ref.[2] |
| **pYFPstar** | | *amyE' cat* TgyrA LIC promoterless *iyfp amyE' bla* ColE1 origin | | Ref.[2] |
| **pRFPstar** | | *amyE' cat* TgyrA LIC promoterless *mCherry amyE' bla* ColE1 origin | | Ref.[3] |
| **pLK** | | *bla spc Pspac comK* P*divIVA lacZ lacI* pLS20-replicon | | Ref.[4] |
| **pDR111** | | *amyE' spc TrrnB* T0λ *Phyperspank*-MCS *lacI amyE' bla* ColE1 origin | | David Rudner Harvard Medical School |
| **pDG1662** | | *amyE' cat amyE' spc bla* | | Ref.[5] |
| **EIB650** | pMAD | pBR322-ori pE194-Ts-ori *rapFphrF*-up/down P*clpB bgaB bla erm* | CQN01/CQN02 + CQN03/CQN04 | This study |
| **EIB188** | pMAD | pBR322-ori pE194-Ts-ori *rapCphrC*-up/down P*clpB bgaB bla erm* | ST74/ST75 + ST68/ST69 | This study |
| **EIB726** | pYFP_bglStar | *bglS'* TgyrA *Phag- iyfp* TgyrA *PtapA-cfp kan bglS'* ColE1 *bla* f1(+) | restriction-enzyme ligation cloning | This study |
| **EIB750** | pDR111 | *amyE' spc TrrnB* T0λ *Phyperspank-lytABC lacI amyE' bla* ColE1 origin | CQN045/CQN047 | This study |
| **EIB751** | pDR111 | *amyE' spc TrrnB* T0λ *Phyperspank-lytC lacI amyE' bla* ColE1 origin | CQN046/CQN047 | This study |
| **EIB752** | pRFPstar | *amyE' cat* TgyrA *PlytABC-mcherry amyE' bla* ColE1 | CQN048/CQN049 | This study |
| **EIB281** | pDG1662 | *amyE' cat Phyperspank_PcomA comA lacI amyE' spc bla* | restriction-enzyme ligation cloning | This study |
| **EIB517** | pDG1662 | *amyE' cat Phyperspank_PcomA comAD55A lacI amyE' spc bla* | MN42/MN43 | This study |

| | | | | |
|---|---|---|---|---|
| **EIB672** | pMAD | pBR322-ori pE194-Ts-ori *lytRABC* -up/down P*clpB bgaB bla erm* | Bu001/Bu002+Bu003/Bu004 | This study |
| **EIB727** | PCFPbgl_Star | *bglS'* TgyrA LIC P$_{tapA}$-*CFP*$_{bs}$ *kan 'bglS* ColE1 *bla* f1(+) | BS0064/BS0065 | This Study |
| **EIB799** | pRFP_Star | P*tapA-lytABC-mCherry* | BS0064/BS0065 +BS0223/BS0225 | This study |

**Supplementary Table S3. Oligonucleotides used in this work.**

| Primer | Sequence 5' – 3' (Restriction cut-site in bold) | Purpose |
|---|---|---|
| **DW120** | AAAGTTGTTGACTTTATCTAC | Verify pLK plasmid |
| **DW121** | CCTCTGATAGACAGCATGTC | |
| **Seq Son 14** | CTGATGGTCGTCATCTACCTGCC | Verify pMAD plasmid |
| **ST126** | ATCATTTATCAACTCTTTTACAC | |
| **CK106** | GCTCGCCATGACTTCACTAAC | *amyE* locus PCR |
| **CK129** | GAGCATTTGCGCTGCTTG | |
| **ST129** | ATGAGTATTCAACATTTCCGTGTC | Verify *amp* gene |
| **ST130** | TTACCAATGCTTAATCAGTGAGG | |
| **DW92** | GTGATGACCATTGCGGCGCTTTTG | *bglS* locus PCR |
| **LA38** | CGGCTCTACAAAGACGAATTTG | |
| **ST 74** | AAA**GTCGAC**AGGGTTGAACAGCCATACGATTCC | Up-fragment of *rapCphrC* |
| **ST 75** | GAGAAGGGGCTTGTTCTTACCTCTTCATTCCTTCACCCTCTCC | |
| **ST 68** | GTAAGAACAAGCCCCTTCTCATTAG | Down-fragment of *rapCphrC* |
| **ST 69** | CAT**AGATCT**CGAAGAGGATTTGCATGCCGATG | |
| **CQN01** | ATGT**GGATCC**GAAGAGCAATCGTTGTCACAAC | Up-fragment of *rapFphrF* |
| **CQN02** | ATCATTCCATCAACATTCTTGTGAGAAGCG | |
| **CQN03** | GTTGATGGAATGATTTAACCGCCGTCC | Down-fragment of *rapFphrF* |
| **CQN04** | ATAGA**GTCGAC**CATCACGTCTAAAGAACTGCTTG | |
| **CQN045** | ACGC**GTCGAC**AAGGAGGAACTACTATGAAAAAATTTATTGCTTTACTG | Amplification of *lytABC* from BIB224 genome |
| **CQN047** | ACAT**GCATGC**TTATCTGTAATAAGATACTGTGC | |
| **CQN046** | ACGC**GTCGAC**AAGGAGGAACTACTATGCGTTCTTATATAAAAGTC | Amplification of *lytC* from BIB224 genome |
| **CQN047** | ACAT**GCATGC**TTATCTGTAATAAGATACTGTGC | |
| **CQN048** | GTTCCTCCTTCCCACCATATTTTATCGTCAACCTATTTTATATTTTAAAG | Amplification of P*lytABC* from BIB224 genome |
| **CQN049** | CCGCGGGCTTTCCCAGCCTGACCATGTCGGTTGTATTC | |
| **Bu001** | TACGCG**GTCGAC**GTCCACTTTGCACCGCTCATGA | Up-fragment of *lytABC* |
| **Bu002** | GTGCGAAGCTTCCATGACGGCACAGTATCTTATTA | |
| **Bu003** | GTGCCGTCATGGAAGCTTCGCACAATGTGCAAAG | Down-fragment of *lytABC* |
| **Bu004** | ACATG**CCATGG**GATACGACGACGACGTTTGC | |
| **MN42** | GCAAGCGGTCCATTGAATACAGCTTGGCATCGATTTTC | site-directed mutagenesis of *comA* |
| **MN43** | CCGACATTCAGCTTATTGAAAATCGATGCCAAGCTG | |

| | | |
|---|---|---|
| **BS0064** | CCGCGGGCTTTCCCAGCCTCAGAGTTAAATGGTATTGCTTCACT | Amplification of PtapA-RBS from EIB727 |
| **BS0065** | GTTCCTCCTTCCCACCTGTAAAACACTGTAACTTGATAT | |
| **BS0223** | GGGAAGGAGGAACTACTATGAAAAAATTTATTGCTTTACTGTTCT | Amplification of lytABC from BIB224 genome |
| **BS0225** | GTTCCTCCTTCCCACCTTATCTGTAATAAGATACTGTGCCGTCA | |

### SUPPLEMENTARY METHODS

#### Compiling of Supplementary Movies

All supplementary movies were compiled using the Fiji software package[6]. Movies were registered with the Linear Stack alignment with SIFT  Plugin[7] using the default settings (rigid transformation, 1.6 px initial gaussian blur, 3 steps per scale, feature descriptor size of  4, 8 feature descriptor orientation bins, closest/nest closest ratio 0.92, maximal alignment error 25 px, inlier ratio 0.05, interpolated output).

#### Fitting of lysis data from QS mutants

Initial lysis rates were determined by analyzing OD curves in initial phases of lysis (50min to 100min). The curves of all mutant strains were fit to satisfy the equation $\mathrm{OD(t)} = \mathrm{f}_i \cdot \mathrm{e}^{-\lambda_1 \mathrm{t}} + (1 - \mathrm{f}_i) \cdot \mathrm{e}^{-\lambda_2 \mathrm{t}}$ with $\mathrm{f}_i$ as the fraction of fast-lysing cells in the *i-th* mutant strain and $\lambda_{1/2}$ as the lysis rates of the two subpopulations. All parameters were determined using fitting using Matlab 2017a (MathWorks Inc.). The model equation was solved numerically using the *ode15s* solver. Parameters were optimized globally by minimizing the sum of squared residuals (SSR) of all data sets with the fsolve function and Levenberg-Marquard function and Levenberg-Marquard algorithm. The confidence intervals (CI) of the resulting solution were determined by fitting to a random subset of 5 techincal replicates for each mutant strain, which was repeated 200 times. The 95% CI was then determined as the 0.025 and 0.975 quantiles of the prediction.

### SUPPLEMENTARY FIGURES

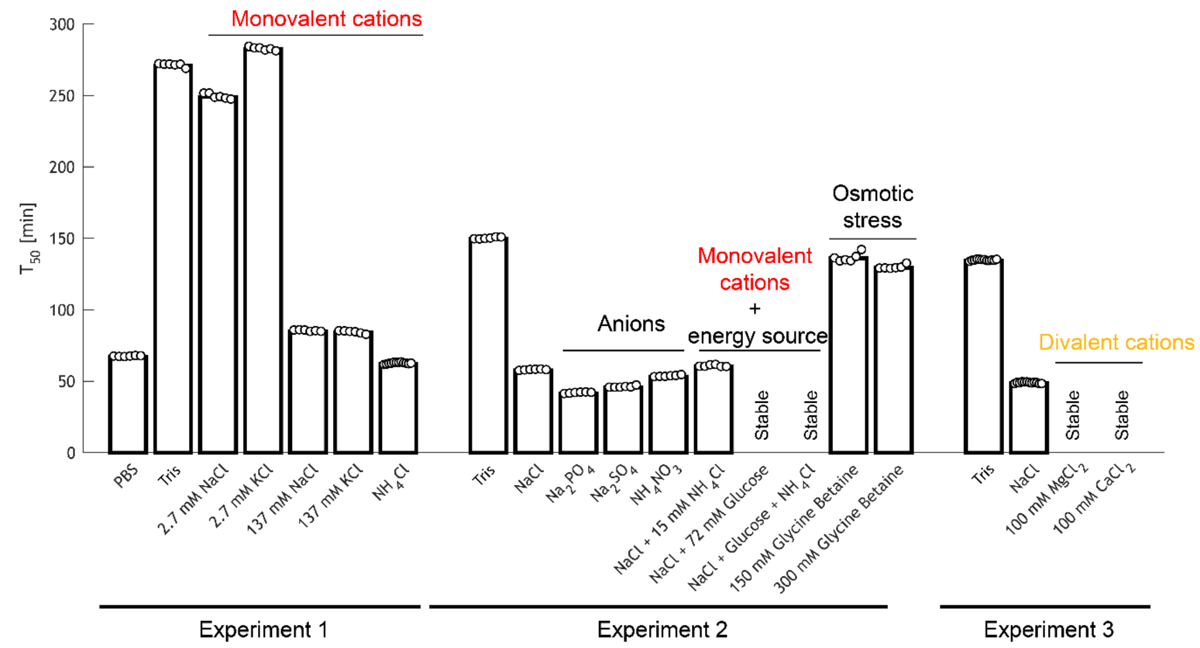


**Supplementary Figure 1. A combination of monovalent ions and nutrient deprivation is lethal for hyposocial *B. subtilis*.** Decline of the optical density $OD_{600nm}$ as a function of time was measured in 96-well plates to determine the exposure time $T_{50}$ that is required to reduce initial OD by 50%. Conditions where OD decline was less than 50% are denoted as NA. Data from different experiments are grouped. Each dot represents results from a single technical replicate.  Substances were added to a final concentration of 137 mM to Tris buffer unless noted otherwise. Glucose was always added at 72 mM. In combination with glucose, $NH_4Cl$ was added at 15 mM.  Addition of glucose or divalent cations prevent lysis. Monovalent cations ($Na^+$, $K^+$, $NH_4^+$) induce lysis regardless of the anion. Addition of glycine betaine has no effect of lysis in Tris-HCl. Strains: BIB1933 (Experiment 1) and BIB2153 otherwise.

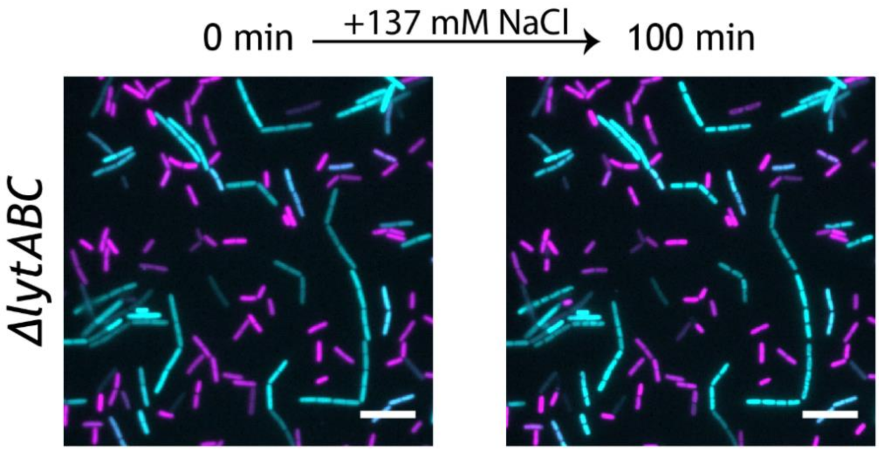


**Supplementary Figure 2. *ΔlytABC* retains both cell-types and does not lyse upon exposure to NaCl.** Micrographs of a *ΔlytABC* before (left) and after exposure to NaCl (right). Both strains carry a reporter for motile ($P_{hag}$-yfp) and sessile cell state ($P_{tapA}$-CFP). Images show superposition of the YFP (magenta) and CFP channel (cyan). Strain: BIB2232. Scale bar: 10 µm.

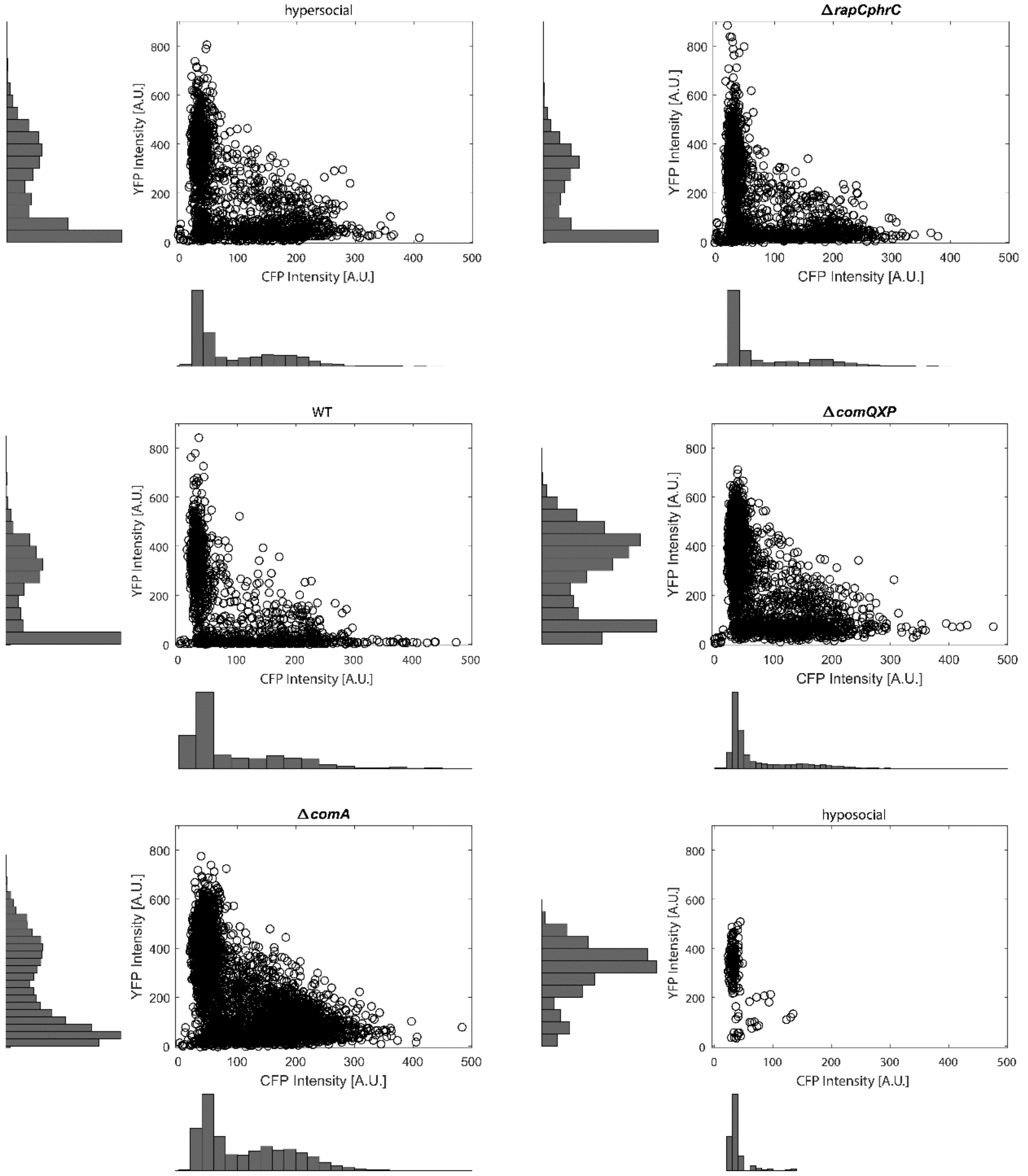


**Supplementary Figure 3. Fluorescence from $P_{hag}$-YFP and $P_{tapA}$-CFP provide a stable read-out to characterize population composition in different mutant backgrounds.** Circles denote the average cellular CFP and YFP intensities as measured in individual cells transformed with the cell-type reporter ($P_{hag}$-YFP $P_{tapA}$-CFP) in the indicated genetic background. Strains from left to right, top-down: BIB2107, BIB2129, BIB2127, BIB2135, BIB2137, BIB2109. The range of fluorescence intensities for YFP and CFP is relatively stable and unaffected by the genotype. Note that the projected fluorescence distributions are not representative for population compositions as sampling of subpopulations on agarose pads was uneven.

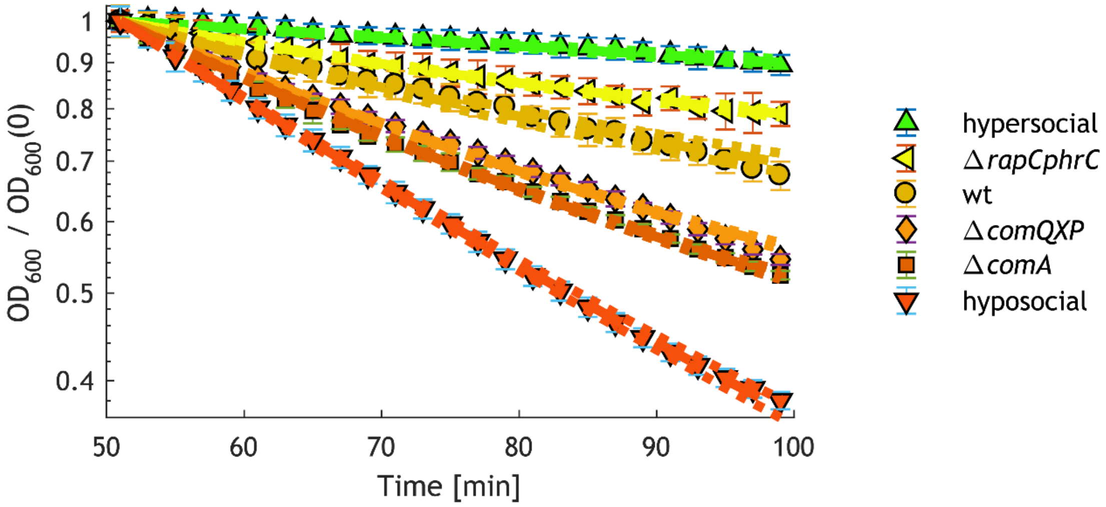


**Supplementary Figure 4. Signaling mutants affect population-level lysis by shifting population composition.** Decline of $OD_{600}$ as a function of time from different regulatory mutants exposed to PBS. Symbols denote experimental data represented as mean ± std for $n_r$ = 8. Lines denote mean and 95% of all model predictions from bootstrapping as obtained from a mathematical model that assumes that regulatory mutations affect solely population composition without changing the lysis rates of individual subpopulations. See Supplementary Methods for details.

### SUPPLEMENTARY MOVIES

**Supplementary Movie S1** (https://doi.org/10.5446/51550). **Exposure to NaCl selectively eliminates the motile subpopulation of cells.** Motile cells express YFP and are false colored in magenta. Sessile cells express CFP and are false colored in cyan. Cells are immobilized on a Tris agarose pad. Upon exposure to NaCl motile cells lyse. Movie shows superimposed brightfield and fluorescence micrographs before adding NaCl, followed by brightfield images only for the duration of lysis. Strain: BIB2127 *bglS::* $P_{tapA}$-*CFP* $T_{gyrA}$ $P_{hag}$-*YFP.* Scale bar: 10 µm.

**Supplementary Movie S2** (https://doi.org/10.5446/51551). **NaCl preferentially lyses LytABC expressing sessile cells.** Cells engineered to express *lytABC* from $P_{tapA}$ (differentially activated in sessile cells) were exposed to NaCl. Motile cells express YFP and are false colored in magenta. Sessile cells express CFP and are false colored in cyan. Cells are immobilized on a Tris agarose pad. Upon exposure to NaCl motile cells lyse. Movie shows superimposed brightfield and fluorescence micrographs before adding NaCl, followed by brightfield images only for the duration of lysis. Strain: BIB2356 *bglS::* $P_{tapA}$-*CFP* $T_{gyrA}$ $P_{hag}$-*YFP amyE::*$P_{tapA}$-*lytABC ΔlytABC.* Scale bar: 10 µm.

**Supplementary Movie S3 (**https://doi/org/10.5446/51553**). NaCl will not induce lysis when motile cells do not express LytABC.** The endogenous autolysin lytABC was deleted. Autolysin expression can be controlled by an IPTG-inducible promoter. Cells were left uninduced. Motile cells express YFP and are false colored in magenta. Sessile cells express CFP and are false colored in cyan. Cells are immobilized on a Tris agarose pad. Upon exposure to NaCl motile cells do not lyse and both cell-types recover upon exposure to salt-free LB (nutrient upshift). Movie shows superimposed brightfield and fluorescence micrographs before addition of NaCl, followed by brightfield micrographs only. A substantial fraction of both cell types resumes growth upon exposure to salt-free LB. Strain: BIB2330: *ΔlytABC bglS::* $P_{tapA}$-*CFP* $T_{gyrA}$ $P_{hag}$-*YFP amyE::* $P_{hyperspank}$-*lytABC lacI.* Scale bar: 10 µm.

**Supplementary Movie S4 (**https://doi/org/10.5446/51552**). NaCl lyses both LytABC expressing motile and sessile cells.** All cells were induced with IPTG to express the autolysin complex LytABC from an ectopic locus. The endogenous autolysin was deleted. Motile cells express YFP and are false colored in magenta. Sessile cells express CFP and are false colored in cyan. Cells are immobilized on a Tris agarose pad. Upon exposure to NaCl both cell-types lyse. Only a small fraction of both cell types resumes growth upon exposure to salt-free LB (nutrient upshift). Movie shows superimposed brightfield and fluorescence micrographs before addition of NaCl, followed by brightfield micrographs only. Strain: BIB2330: *ΔlytABC bglS::* $P_{tapA}$-*CFP* $T_{gyrA}$ $P_{hag}$-*YFP amyE::* $P_{hyperspank}$-*lytABC lacI.* Scale bar: 10 µm.

**Supplementary Movie S5 (**https://doi/org/10.5446/5154**). Exposure to KCl selectively eliminates the motile subpopulation of cells.** Motile cells express YFP and are false colored in magenta. Sessile cells express CFP and are false colored in cyan. Cells are immobilized on a Tris agarose pad. Upon exposure to KCl motile cells lyse. Movie shows superimposed brightfield and fluorescence micrographs before addition of NaCl, followed by brightfield micrographs only. Strain: BIB2246 *bglS::* $P_{tapA}$-*CFP* $T_{gyrA}$ $P_{hag}$-*YFP amyE::*$P_{lytABC}$-*mCherry.* Scale bar: 10 µm.

**Supplementary Movie S6 (**https://doi/org/10.5446/51555**). Exposure to NaCl selectively eliminates the motile subpopulation of cells of an undomesticated soil isolate PS216.** Motile cells express YFP and are false colored in magenta. Sessile cells express CFP and are false colored in cyan. Cells are immobilized on a Tris agarose pad. Upon exposure to NaCl motile cells lyse. Movie shows superimposed brightfield and fluorescence micrographs. Strain: BIB2051: *amyE::* $P_{tapA}$-*CFP* $T_{gyrA}$ $P_{hag}$-*YFP.* Scale bar: 10 µm.
